# Eucalyptus microRNA Archive (EMA): a multi-study and cross-condition curated database of microRNAs in *Eucalyptus grandis*

**DOI:** 10.64898/2026.08.29.747619

**Authors:** João Vítor Aires-Teixeira, Thiago Motta Venancio, Gabriel Quintanilha-Peixoto, Kellen Kauanne Pimenta de Oliveira

## Abstract

MicroRNAs (miRNAs) are key post-transcriptional regulators of development, stress response, and secondary cell wall formation in woody plants, yet annotations for *Eucalyptus grandis*, the world’s most widely planted hardwood, remain fragmented across studies using incompatible discovery pipelines and filtering criteria. Here we present the Eucalyptus MicroRNA Archive (EMA), a curated, locus-resolved database integrating three independent small RNA sequencing datasets spanning vegetative tissue, somatic embryogenesis, and mechanically induced tension wood formation. Applying annotation criteria aligned with current plant miRNA standards, EMA catalogs 99 curated miRNAs (31 previously described, 68 novel) organized into 34 family-level groupings under a three-tier confidence system, known-reference-supported, multi-study replicated, or single-study, that preserves study-of-origin and sample-level evidence for every entry. Cross-study comparison showed that only 9 of 99 entries (9.1%) were independently supported by all three datasets, supporting an evidence-tiered rather than binary annotation scheme. Target prediction against the *E. grandis* transcriptome yielded 1,773 miRNA-target interactions spanning 764 loci, integrated into a combined miRNA-target and protein-protein interaction network. This network resolved into functionally coherent, mutually isolated clusters, including an miR482-associated NBS-LRR/TIR disease-resistance hub with a substantial translational-repression component, alongside modules enriched for ribosome biogenesis and translation, DNA replication, and nitrogen and carbohydrate metabolism. EMA is publicly accessible through an interactive web dashboard, with all curated data, source code, and analysis scripts openly available, providing a reproducible, extensible framework for *E. grandis* miRNA research and a template for similarly structured resources in other non-model woody species.

## 1 INTRODUCTION

MicroRNAs are small non-coding RNAs that act as post-transcriptional regulators of gene expression, generally through sequence-guided transcript cleavage or translational repression. In plants, these molecules are commonly associated with development stages, organ differentiation, hormone signaling, stress responses, and environmental adaptation [1]. Their biological relevance is particularly important in long-lived woody species, where the timing of the juvenile-to-adult phase transition, the differentiation of xylem and phloem during radial growth, and the deposition of lignin and cellulose in the secondary cell wall all depend on tightly timed, tissue-specific gene expression, a regulatory precision to which miRNA-mediated post-transcriptional control is particularly well suited [2,3].

In *Eucalyptus grandis*, a reference species for forest biotechnology and woody plant genomics, miRNAs have been reported in several studies since 2014, in contexts ranging from genome annotation to vegetative development, somatic embryogenesis and tension wood formation [4–9]. These publications differ in sequencing platform, tissue, filtering criteria, and reported family counts, while their miRNA annotations remain distributed across independent analytical frameworks with limited consolidation between them.

The rapid expansion of small RNA sequencing has increased the number of predicted plant miRNAs, but it has also intensified concerns about annotation consistency and false positives; plant genomes produce various classes of small RNAs, and it is debatable whether every hairpin-associated or abundant small RNA should be interpreted as a *bona fide* miRNA [10,11]. Updated plant miRNA annotation criteria emphasize the importance of precursor structure, precise processing, miRNA star evidence, reproducibility, and careful exclusion of other non-coding RNA classes. Broad plant databases such as PmiREN [12] have addressed these issues by applying standardized processing, strict annotation criteria, and comprehensive annotation across multiple species. PmiREN 3.0 currently lists 82 miRNA loci in 58 families and 6 clusters for *E. grandis,* with no associated sRNA-seq or PARE-seq datasets. Species-focused resources remain valuable when they retain study-level and condition-specific information that may be absent from broader multispecies summaries [13].

In this study, we present the Eucalyptus MicroRNA Archive (EMA), a curated database and online resource for miRNA discovery, annotation, and functional prioritization in *E. grandis*. EMA integrates public small RNA sequencing datasets from three independent studies while preserving study-specific evidence through a conservative locus-level union strategy. In the current release, EMA v1.0 contains 99 curated miRNAs, including 31 previously described and 68 novel candidates. The curated miRNAs are organized into 34 family-level groupings, 12 of which correspond to families previously represented by known miRNAs in *E. grandis.* Each miRNA is associated with precursor features, expression profiles, differential expression evidence, and predicted targets, which were further integrated into a miRNA–target and protein–protein interaction network. By combining miRNA discovery, a conservative post-processing filtering aligned with plant miRNA annotation criteria, and a relational database implementation, EMA provides a reproducible framework for exploring conserved and newly curated miRNA candidates in *E. grandis*.

## 2 METHODS

All miRNAs incorporated into the final database were curated under an evidence-conservative framework guided by updated annotation criteria for plants proposed by Axtell and Meyers (2018) [1], revising the original 2008 community standard [14] to curb false-positive annotations in the deep-sequencing era.

### 2.1 Public small RNA sequencing datasets and study design

Dataset selection followed strict inclusion criteria to ensure biological interpretability and technical comparability: (i) *E. grandis* as the model species; (ii) well-defined biological contexts, developmental or physiological; (iii) availability of raw sequencing data in public repositories; and (iv) sufficient sequencing depth.

A total of three independent studies comprising 20 samples were selected, covering complementary biological contexts including vegetative tissues, somatic embryogenesis-related tissues and mechanically induced wood formation [7–9]. miRNA discovery and genomic coordinate assignment were performed using the *E. grandis* Phytozome v2 reference genome [15].

**Table 1.**
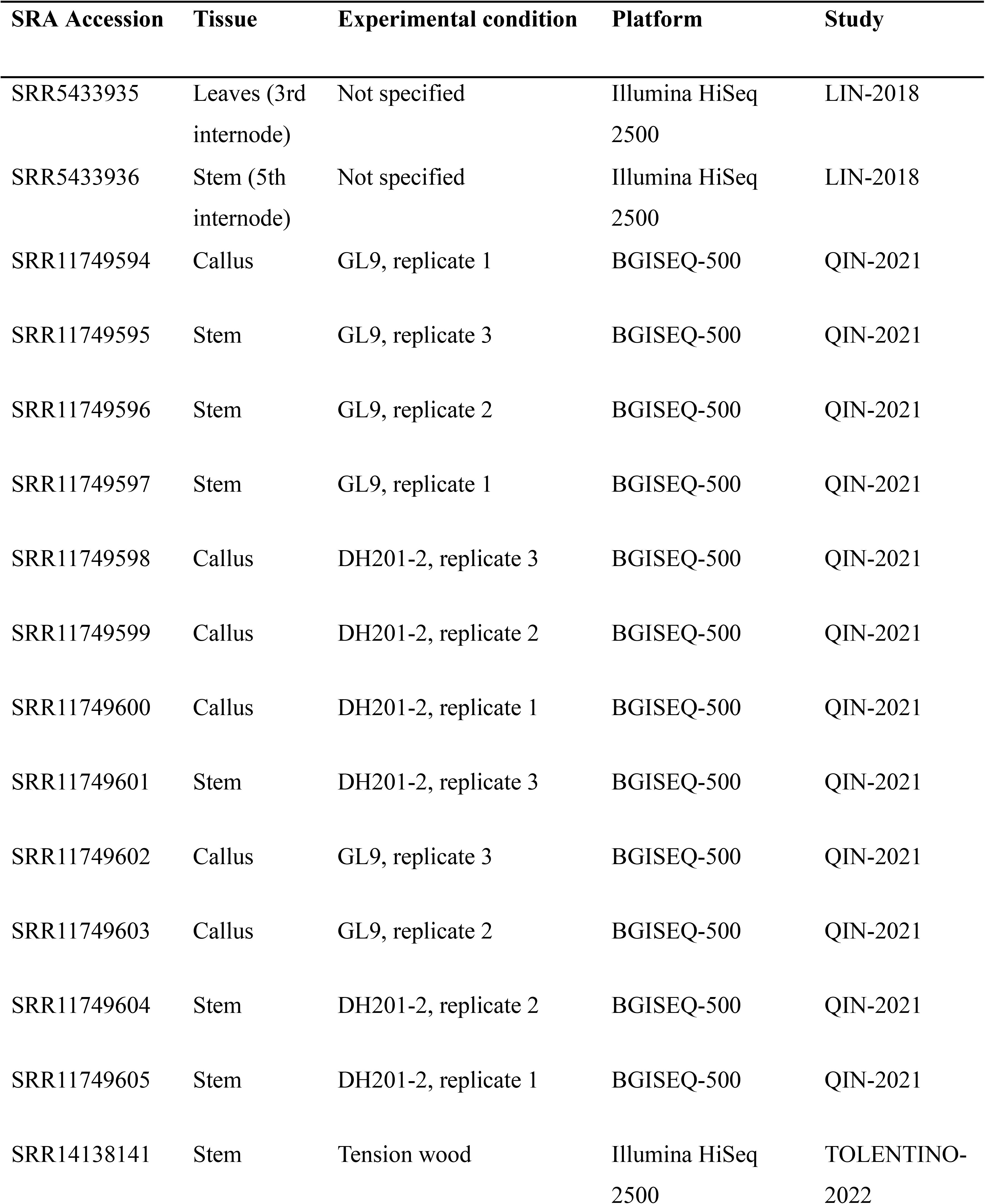
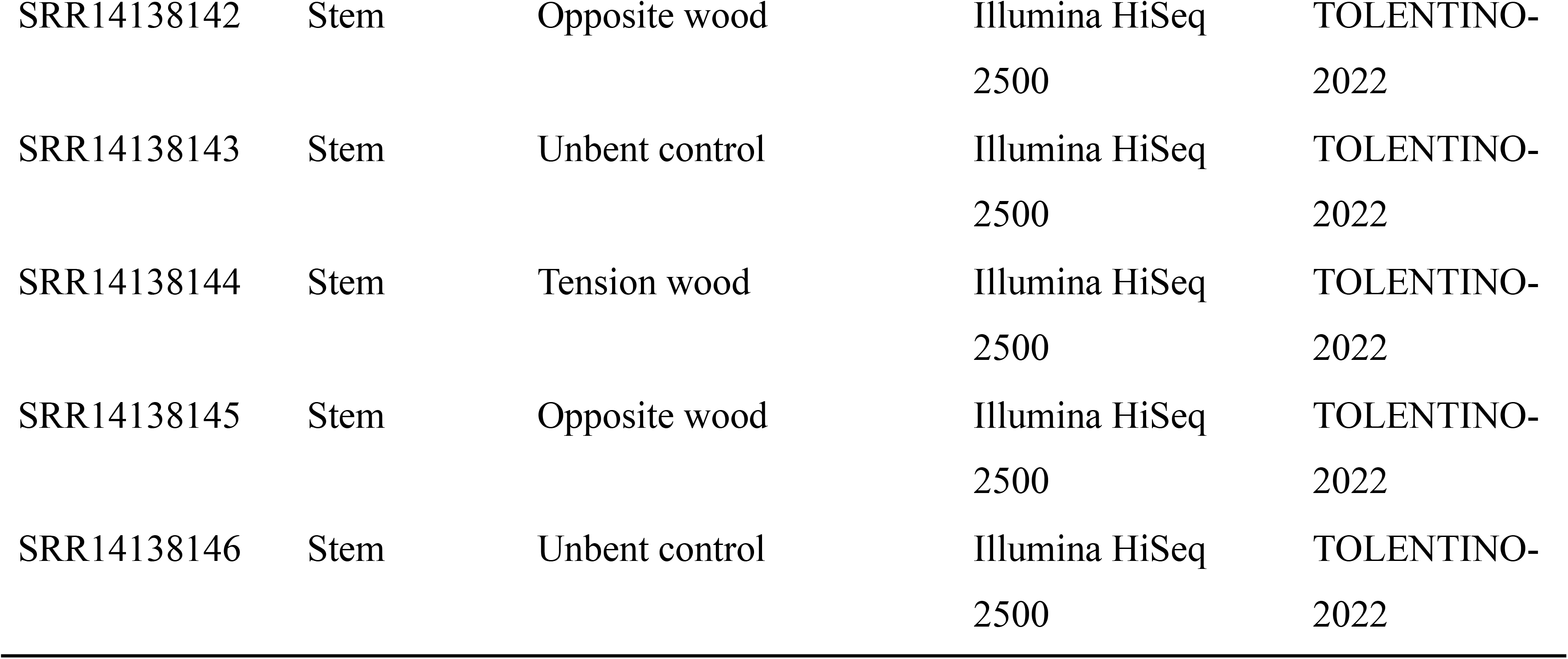
Summary of samples used for miRNA discovery and analysis.

| **SRA Accession** | **Tissue** | **Experimental condition** | **Platform** | **Study** |
| --- | --- | --- | --- | --- |
| SRR5433935 | Leaves (3rd internode) | Not specified | Illumina HiSeq 2500 | LIN-2018 |
| SRR5433936 | Stem (5th internode) | Not specified | Illumina HiSeq 2500 | LIN-2018 |
| SRR11749594 | Callus | GL9, replicate 1 | BGISEQ-500 | QIN-2021 |
| SRR11749595 | Stem | GL9, replicate 3 | BGISEQ-500 | QIN-2021 |
| SRR11749596 | Stem | GL9, replicate 2 | BGISEQ-500 | QIN-2021 |
| SRR11749597 | Stem | GL9, replicate 1 | BGISEQ-500 | QIN-2021 |
| SRR11749598 | Callus | DH201-2, replicate 3 | BGISEQ-500 | QIN-2021 |
| SRR11749599 | Callus | DH201-2, replicate 2 | BGISEQ-500 | QIN-2021 |
| SRR11749600 | Callus | DH201-2, replicate 1 | BGISEQ-500 | QIN-2021 |
| SRR11749601 | Stem | DH201-2, replicate 3 | BGISEQ-500 | QIN-2021 |
| SRR11749602 | Callus | GL9, replicate 3 | BGISEQ-500 | QIN-2021 |
| SRR11749603 | Callus | GL9, replicate 2 | BGISEQ-500 | QIN-2021 |
| SRR11749604 | Stem | DH201-2, replicate 2 | BGISEQ-500 | QIN-2021 |
| SRR11749605 | Stem | DH201-2, replicate 1 | BGISEQ-500 | QIN-2021 |
| SRR14138141 | Stem | Tension wood | Illumina HiSeq 2500 | TOLENTINO-2022 |
| SRR14138142 | Stem | Opposite wood | Illumina HiSeq<br>2500 | TOLENTINO-2022 |
| SRR14138143 | Stem | Unbent control | Illumina HiSeq<br>2500 | TOLENTINO-2022 |
| SRR14138144 | Stem | Tension wood | Illumina HiSeq<br>2500 | TOLENTINO-2022 |
| SRR14138145 | Stem | Opposite wood | Illumina HiSeq<br>2500 | TOLENTINO-2022 |
| SRR14138146 | Stem | Unbent control | Illumina HiSeq<br>2500 | TOLENTINO-2022 |

For LIN-2018 samples, leaf (third internode) and stem (fifth internode) tissue were sampled from 5-month-old *E. grandis* trees under standard growth conditions, with no experimental treatment applied beyond normal vegetative development; each tissue library was pooled from three individual trees prior to sequencing [7]. QIN-2021 sampled differentiated stem tissue and dedifferentiated, tissue-culture-induced callus (10 days on 2,4-D-supplemented induction medium) from two genotypes, GL9 and DH201-2 (an *E. grandis × urophylla hybrid),* with three biological replicates per tissue-genotype combination, to capture miRNA dynamics associated with differences in somatic embryogenic potential between the two genotypes [8]. TOLENTINO-2022 induced reaction wood by mechanically bending *E. grandis* stems at a 45° angle for one month; bent stems were longitudinally bisected into tension wood (TW, outer face) and opposite wood (OW, inner face) segments and sampled alongside unbent control (CT) stems from independent, unrelated trees, with two biological replicates per condition [9].

### 2.2 Quality control of sequencing reads

Raw reads were processed with fastp v1.0.1 [16]. Reads were retained within an 18–24 nt size window and required to meet a minimum Phred quality score of 20, with no more than 40% low-quality bases per read and no more than five ambiguous (N) bases. Adapter and 3’ quality trimming used a four-base sliding window with a minimum mean quality of 20. Poly-G artifacts were removed using a 10 nt minimum run-length threshold, and low-complexity reads were excluded at a complexity threshold of 30%. Duplication-rate estimation was disabled, as high redundancy of identical reads is expected and biologically meaningful in small RNA libraries [17].

### 2.3 miRNA discovery

miRNA discovery was performed independently for each study using miRDeep2 v2.0.1.3 [18]. We used miRDeep2 as the main discovery software due to its read-collapsing, genome-mapping, and hairpin-scoring framework [19]. miRDeep2 outputs were subjected to a dedicated post-processing filtering layer aligned with the plant miRNA annotation criteria of Axtell and Meyers (2018).

For each study, miRDeep2 result tables and output files were parsed to extract mature sequence, star sequence, precursor sequence, precursor coordinate, miRDeep2 score, read count metrics, randfold status, RFAM alert, conservation seed score, and true positive probability.

### 2.4 Candidate miRNA curation and classification

Known miRNAs were retained when a valid reference miRNA identifier was present, and the miRDeep2 discovery score was at least 10. Similarity-based identification of known miRNAs relied on a combined reference set of mature and hairpin sequences, comprising entries specific to *E. grandis* from PmiREN v2.0 [12] and the full *Viridiplantae* collection from miRBase [20]. This threshold was chosen as a conservative cutoff considering that the discovery scores varied by several orders of magnitude across the three studies: per-study mean scores ranged from 9,682.2 (LIN-2018) to 140,709.8 (TOLENTINO-2022), while individual scores across all evidence records spanned from 0.4 to 3,724,090.8 overall. The value 10.0 was selected to exclude the lowest-confidence tail of candidate detections while retaining the substantial majority of genuine known-miRNA signals. Novel candidates were subjected to a more strict filtering layer adapted from the plant miRNA annotation recommendations of Axtell and Meyers (2018), applied to candidates lacking a reference miRNA identifier and implemented on top of the structural and read-signature evidence produced by miRDeep2. Applying the novel-candidate filtering criteria (mature length restriction, star-read minimum, precursor length ceiling) to known miRNAs would have incorrectly discarded legitimate known entries whose mature sequence falls outside the 20 to 22 nt window used for novel candidates (observed known mature lengths ranged from 19 to 23 nt).

A novel candidate was retained only when it met all of the following conditions, adapted from the plant miRNA annotation criteria of Axtell and Meyers (2018), that is, absence of an RFAM alert; randfold p-value equal to or lower than 0.05, as computed internally by miRDeep2 using the approach of [21]; at least 10 star reads; precursor length no greater than 300 nt; mature length between 20 and 22 nt; and operational processing precision of at least 0.75, corresponding to the minimum threshold recommended by those criteria. Processing precision was calculated as:

*Precision = (mature read count + star read count) / total read count*

Family-level assignment for both known and novel candidates was derived from the miRDeep2 seed-matching output field, which compares the seed region (nucleotides 2 to 8) of each candidate mature sequence against the full miRBase reference set across species. This mechanism allows conserved plant families to be recognized even when the candidate itself has never been annotated in *E. grandis*.

### 2.5 Locus-level union integration and cross-study evidence reconciliation

miRNA candidates were reconciled using a conservative locus-level union strategy. This approach was adopted because the source studies differed in biological context, sequencing platform, sample structure, and sequencing depth; a strict intersection across all studies would be overly restrictive and could exclude context-dependent miRNAs.

For novel candidates, precursor loci were grouped when they occurred on the same chromosome or scaffold, on the same strand, and showed at least 80 percent reciprocal precursor overlap; the candidate with the highest miRDeep2 score within each cluster was selected as the canonical representative. Known miRNAs were integrated using their reference-supported miRNA identifiers rather than locus overlap alone; when the same known miRNA was detected in more than one study, the highest-scoring record was selected as the canonical representative.

All additional study-specific observations were retained in an evidence table and classified according to their relationship to the canonical entry: direct canonical evidence (the record selected as representative), same-mature-sequence evidence (an identical mature product observed at another locus or in another study), same-identifier evidence (a record independently assigned the same reference miRNA name), or same-locus-overlap evidence (a genomically overlapping observation on the same strand).

To prevent cross-attribution between distinct known miRNAs that happen to share an identical mature sequence, reconciliation by shared mature sequence was applied only when the observed sequence corresponded to a single known miRNA identifier in the catalog; observations whose mature sequence matched more than one distinct known entry, or whose independently miRDeep2-assigned identifier explicitly named a different miRNA, were excluded from the evidence set of any single entry rather than attributed by sequence coincidence.

Because standard short-read quantification tools cannot distinguish reads originating from genomically distinct loci that produce an identical mature sequence [22,23], all catalog entries were screened for shared mature sequences. Exact sequence identity and single-nucleotide 5’ or 3’ terminal variation were the two criteria, reflecting the well-documented imprecision of Dicer/DCL1 cleavage at hairpin termini rather than an arbitrary edit-distance threshold [24]. Transitive closure was applied across these pairwise relationships (connected-component grouping), so that loci differing by more than one nucleotide but linked through a shared intermediate variant were merged into a single group.

Expression and differential expression values are identical among members of the same group; each member retains an independent genomic locus, precursor structure, and, where applicable, independent discovery evidence, and grouping does not affect miRNA discovery, precursor curation, or target prediction, which remain locus-specific. It is critical to distinguish discovery from expression: while locus-specific discovery evidence strictly excluded ambiguous mature matches, expression quantification was inherently collapsed by mature sequence groups, reflecting the biological reality of Dicer processing [1].

### 2.6 Expression quantification

Expression quantification was evaluated using the quantifier module of miRDeep2 in all 20 samples, using a reference of mature and precursor sequences built exclusively from the curated catalog entries. Counts per million were calculated using the total number of reads mapped to the *E. grandis* reference genome for each sample as the library-size denominator, avoiding circular normalization against the miRNA reference itself [25].

### 2.7 Differential expression analysis

Differential expression was performed on the raw counts with DESeq2 v1.48.1 [26]. miRNAs were retained for testing when they had at least 10 raw counts in at least two samples within each study subset. QIN-2021 [8] was analyzed using a group-based design and a false discovery rate threshold of 0.05, across four pairwise contrasts (GL9 stem vs. GL9 callus, DH201-2 stem vs. DH201-2 callus, GL9 callus vs. DH201-2 callus, GL9 stem vs. DH201-2 stem).

For TOLENTINO-2022 [9], tension wood (TW) and opposite wood (OW) samples originated from longitudinally bisected segments of the same bent stem of a given tree, while the unbent control (CT) samples derived from independent, unrelated individuals that were not subjected to bending. Because this asymmetric structure invalidates a single uniform design across all three conditions [27], a mixed statistical approach was applied in order to use TW-vs-OW as a paired model contrast. LIN-2018 [7] was not submitted to differential expression analysis due to the absence of paired replicates per condition in the available samples.

### 2.8 Target prediction

Mature sequences from the curated EMA catalog were submitted to psRNATarget (2017 release, Schema V2) [28] against the reference transcriptome. The expectation (mismatch penalty) threshold was set to 3, and target-site accessibility was constrained to UPE (Unpaired Energy) <= 25. Predicted interactions were stored with miRNA accession, target transcript accession, locus, pacid, annotation version, expectation score, UPE, alignment coordinates, aligned fragment sequences, and inhibition mechanism (cleavage or translational repression). Functional target interpretation used transcript accession, locus name, transcript version, Pfam, Gene Ontology terms, and the best *Arabidopsis thaliana* hits when available.

### 2.9 Network construction and functional analysis

Predicted target loci of the curated miRNA catalog were submitted directly to the STRING database [29] through the stringApp for Cytoscape [30], retrieving physical and functional protein partners in *E. grandis*. The network was built using the full STRING interaction set for *E. grandis*, with a minimum required interaction confidence score of 0.700, no additional interactors beyond the submitted target proteins, and Smart Delimiters enabled for identifier resolution. Nodes without any retrieved interactions were removed from the network, yielding four mutually disconnected components (section 3.6). Each component was manually assigned a descriptive cluster label based on the identity and shared function of its member proteins.

Gene Ontology (GO) Biological Process enrichment for the retrieved protein set was performed through the stringApp enrichment module, using the full *E. grandis* genome as the statistical background. Enrichment results (false discovery rate, gene counts, and background counts per term) were exported from Cytoscape and used to generate a dot plot in R v4.5.1 with the ggplot2 package [31], displaying the 20 most significant terms ranked by false discovery rate, with dot size denoting the number of network proteins annotated to each term and color denoting false discovery rate.

Network visualization and topological statistics calculations were performed using Cytoscape v3.10.4 [30]. Discrete mapping was applied to node attributes to differentiate cluster membership by color.

### 2.10 Database and web interface implementation

The database architecture was developed using SQLAlchemy v2.0.25 [32], a database toolkit for Python v3.13.5. All data was stored in an SQLite database. The backend system was developed using the FastAPI framework v0.109.0 [33], and the RESTful API utilizes Pydantic v2.5.3 [34] for data validation and schema management. The web interface was implemented using the React v19.2.0 framework with TypeScript v5.9.3, powered by the Vite v7.2.4 build tool. It also uses the Bootstrap v5.3.3 framework for styling, alongside SASS, HTML5, and Lucide-React for iconography. EMA v1.0 build is available at https://jvtarss.github.io/ema-dashboard-git.

## 3 RESULTS

### 3.1 Composition of the EMA catalog

The Eucalyptus MicroRNA Archive (EMA) comprises 99 curated miRNAs in *E. grandis*, including 31 previously described (known) and 68 novel candidates. Classified entries span 34 distinct family-level groupings, of which 12 correspond to families represented by at least one previously described miRNA in *E. grandis*; the remaining 22 groupings are populated by novel candidates assigned to conserved plant miRNA families based on seed-sequence similarity. 17 novel candidates, 25.0% of all novel entries, could not be assigned to any recognized family and were unclassified.

The five most represented families, miR156 (13 members, entirely known), miR399 (9 members, entirely novel), miR169 (7 members, entirely novel), and miR167 and miR171 (4 members each), together account for 37 of 99 catalog entries (37.4%). At the other end of the distribution, 18 of 34 families (52.9%) are represented by a single member each, several of them bearing numerical designations in the thousands such as miR5050, miR5139 and miR8717.

### 3.2 Genomic distribution across chromosomes and scaffolds

The genomic distribution of the 99 curated loci across the *E. grandis* chromosomes and unplaced scaffolds is presented in Figure 1.

**Figure 1.**
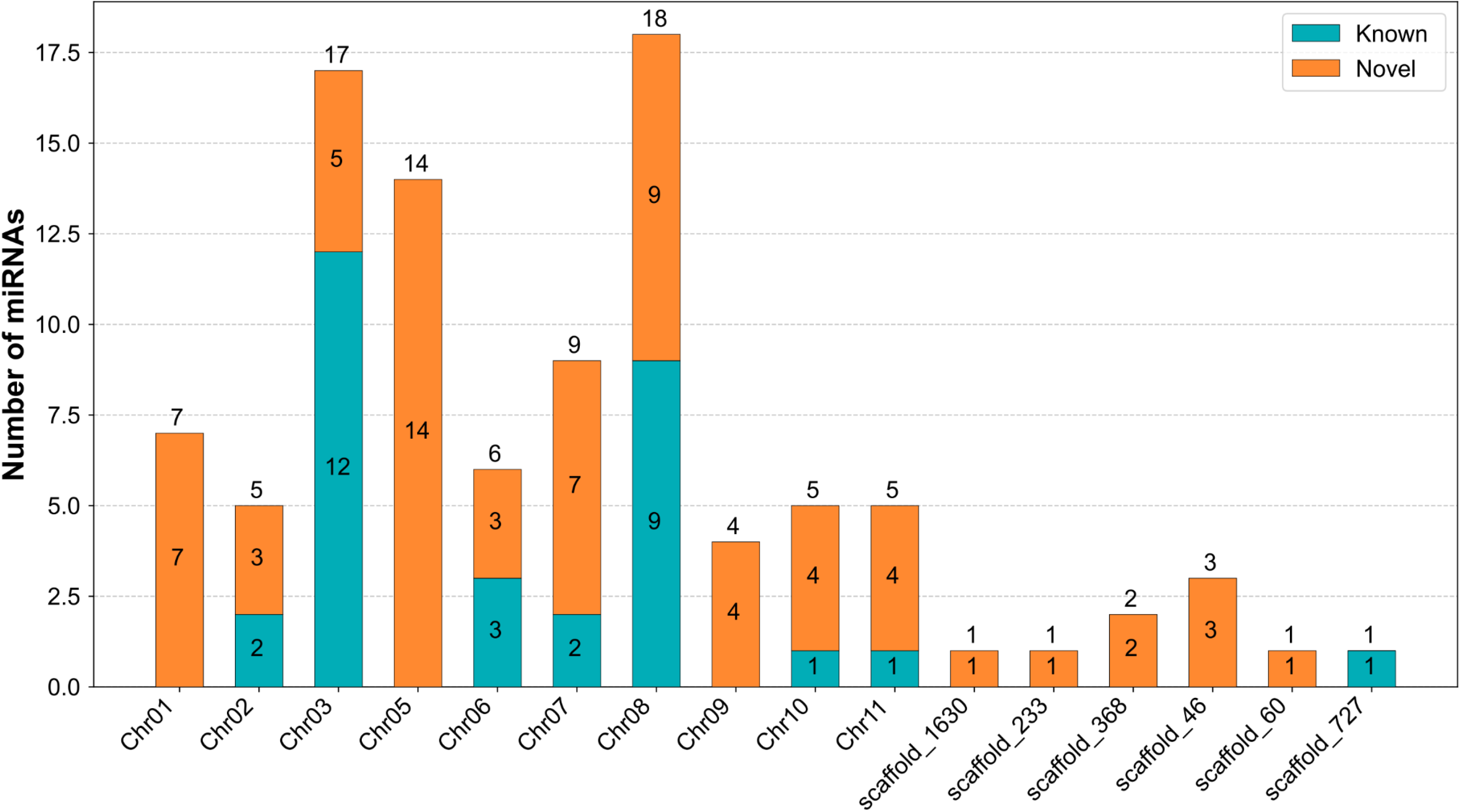
Genomic distribution of main miRNA loci across *E. grandis* chromosomes and scaffolds in the current Phytozome v2 genome assembly.

Known miRNA loci are concentrated on a limited subset of chromosomes: Chr03 alone harbors 12 known loci, the largest known-miRNA cluster in the genome, and Chr08 contributes a further 9 known loci; together these two chromosomes account for 21 of the 31 known miRNAs in the catalog (67.7%) (Figure 1). Novel candidates are distributed more broadly, populating every chromosome as well as six unplaced scaffolds (scaffold_46, scaffold_60, scaffold_233, scaffold_368, scaffold_727 and scaffold_1630). Novel-locus density peaks on Chr05 (14 loci, the largest single novel cluster genome-wide) and Chr01 (7 loci); neither chromosome contains a single previously described miRNA. Chr08 is the only chromosome with an exactly balanced contribution (9 known, 9 novel), whereas Chr03, despite harboring the largest known cluster, contributes comparatively few novel loci (5). The six unplaced scaffolds collectively contribute 9 loci (8 novel, 1 known).

Mature sequence length ranged from 19 to 23 nt among known entries, with an average of 20.45 nt, and was constrained to 20-22 nt among novel entries by the filtering criteria. Precursor lengths (mean 78.6 nt, range 54-90 nt) fall well within the expected range for curated plant stem-loop precursors, and no precursor approached the 300 nt filtering ceiling.

### 3.3 Study-specific evidence and cross-study support

The contribution of each source study to the final catalog, together with patterns of cross-study overlap and discovery-score distributions, is summarized in Figure 2.

**Figure 2.**
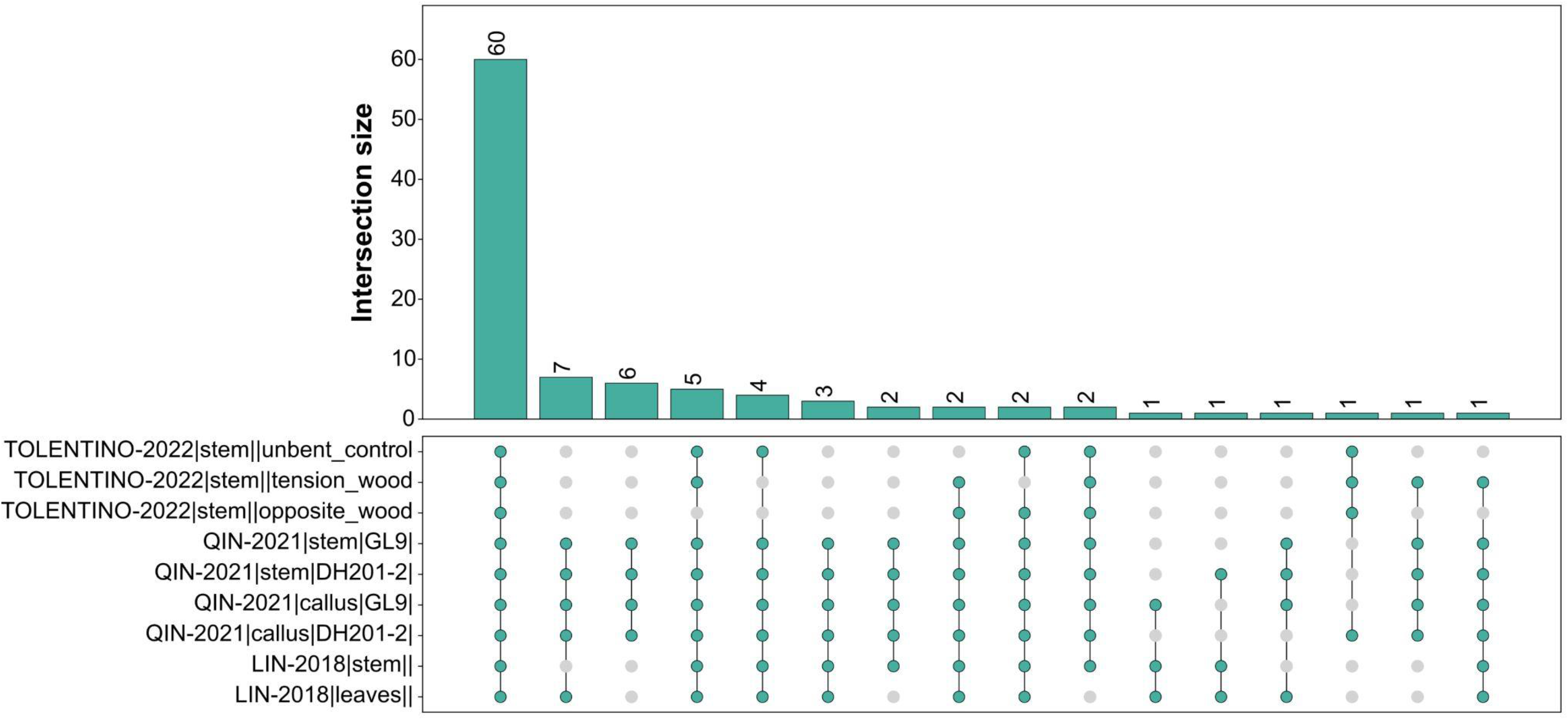
UpSet plot of intersection sizes across the nine sample-group categories spanning the three source studies.

QIN-2021 supported 75 of the 99 catalog entries (75.8%), LIN-2018 supported 44 (44.4%), and TOLENTINO-2022 supported 20 (20.2%) (Figure 2B). The UpSet plot (Figure 2A) shows a single dominant intersection of 60 miRNAs, corresponding to loci detected across all four QIN-2021 sample subgroups (both GL9 and DH201-2 genotypes, in both stem and callus tissue) independently of detection in LIN-2018 or TOLENTINO-2022; all remaining intersections are considerably smaller, ranging from 1 to 7 members, forming a long tail of study- and condition-specific combinations. Consistent with this pattern, 68 of 99 miRNAs (68.7%) are supported by discovery-quality evidence from a single study only, 22 (22.2%) from two studies, and 9 (9.1%) from all three studies.

### 3.4 Structural and read-support features of curated miRNAs

Mature sequence length is concentrated in the 20-21 nt window across the entire catalog (Figure 3A). Known entries span 19-23 nt (mean 20.45 nt), while novel candidates are constrained to 20-22 nt; within this constrained range, the large majority of novel candidates fall at 20 or 21 nt, with comparatively few reaching the upper 22 nt boundary.

**Figure 3.**
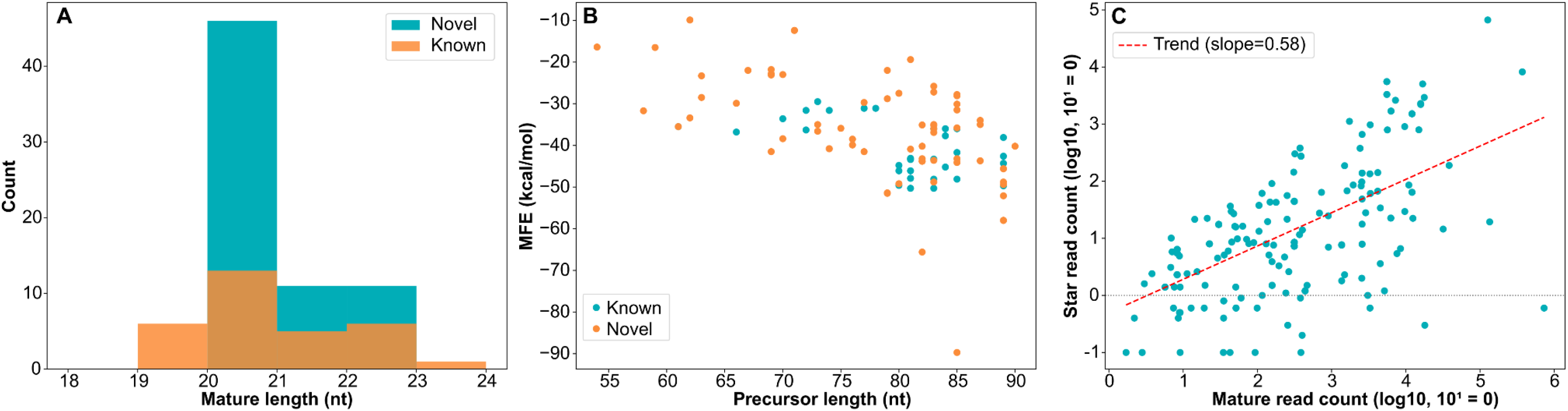
Structural and read-support features of curated EMA miRNAs. (A) Distribution of mature sequence length (nt) for novel and known entries. (B) Precursor length (nt) versus minimum free energy (MFE, kcal/mol), colored by confidence tier: known, reference-supported; novel, multi-study replicated; novel, single-study. (C) Star read count versus mature read count (both log10-transformed), with an ordinary least-squares trend line; the dotted grey line marks zero star reads.

Precursor length and MFE, plotted by confidence tier, are shown in Figure 3B. Known, reference-supported miRNAs cluster at longer precursor lengths (predominantly 70-90 nt) and comparatively stable MFE values (mostly between -30 and -50 kcal/mol). Novel, multi-study replicated candidates occupy a similar but sparser region of this space. Novel, single-study candidates span the entire range of both axes, including several short precursors (54-66 nt) with weak predicted stability (MFE as low as -10 to -20 kcal/mol) and one precursor (∼85 nt) with an unusually low MFE outlier (∼-90 kcal/mol). The dataset-wide mean MFE (-37.76 kcal/mol) and AMFE (-47.70 kcal/mol per nucleotide) summarize this distribution.

Star-read evidence plotted against mature-read abundance (Figure 3C) shows a positive association on the log10 scale (ordinary least-squares slope = 0.58). A subset of loci lies at or near the star-read floor (log10 star count ≈ 0) even at moderate-to-high mature read counts; since the ≥10 star-read requirement was applied only to novel candidates, these floor-level points correspond to known miRNA entries, which are not subject to that criterion.

### 3.5 Expression profiles and differential expression

The unified quantification step, performed against the 99-entry curated reference, produced a raw count matrix covering all 20 samples across the three studies, normalized to counts per million using genome-mapped read totals per sample. Expression patterns across stem-tissue samples from the three studies are shown in Figure 4.

**Figure 4.**
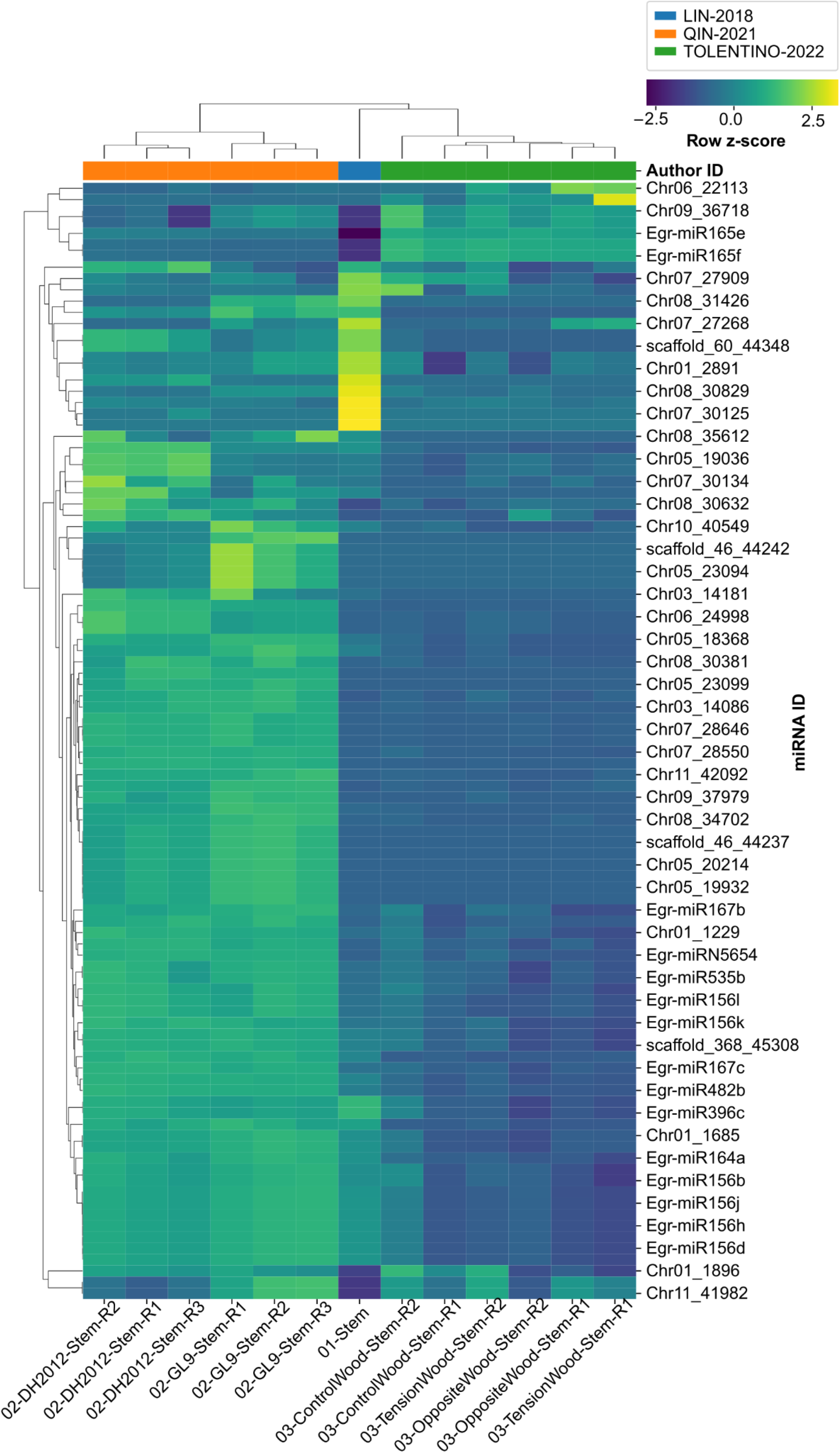
Row-standardized expression heatmap of curated miRNAs across stem-tissue samples from three source studies. Expression values are shown as z-scores, representing the number of standard deviations a sample’s log10(CPM+1) value deviates from that miRNA’s mean expression across all samples. Both samples (columns) and miRNAs (rows) are hierarchically clustered, with column annotations indicating the study of origin. Sample prefixes correspond to: 01 (LIN-2018), 02 (QIN-2021), and 03 (TOLENTINO-2022).

Sample clustering in Figure 4 groups replicates by study of origin in all cases. A small set of miRNAs (Chr07_27909, Chr08_31426, Chr07_27268, scaffold_60_44348, Chr01_2891, Chr08_30829 and Chr07_30125) show markedly elevated z-scores restricted to the single LIN-2018 stem sample, remaining low in both QIN-2021 and TOLENTINO-2022. A larger block of entries, including several miR156 paralogs (Egr-miR156b/d/h/j/k/l), Egr-miR164a, Egr-miR396c and Egr-miR482b, shows consistently elevated expression across QIN-2021 stem replicates and comparatively low expression across all TOLENTINO-2022 stem samples regardless of bending condition. Within TOLENTINO-2022, condition-specific (tension-vs. opposite-vs. control-wood) variation is visually subtle for nearly all miRNAs, with one visible exception (Chr09_36718, elevated in one tension-wood replicate).

Differential expression analysis in QIN-2021 samples (group-based design, FDR < 0.05) tested differences between differentiated stem tissue and dedifferentiated, tissue-culture-induced callus within each of the two genotypes sampled (GL9 and DH201-2), as well as between genotypes within each tissue, given their contrasting somatic embryogenic potential. This identified 223 significant miRNA × comparison instances across the four pairwise contrasts: 78 in GL9 stem vs. GL9 callus, 63 in DH201-2 stem vs. DH201-2 callus, 58 in GL9 stem vs. DH201-2 stem, and 24 in GL9 callus vs. DH201-2 callus. Because 51 of the 99 catalog entries belong to paralogous groups sharing an identical mature sequence and, by construction, an identical expression signal (section 2.6), these raw per-accession counts were re-evaluated at the level of distinct expression signals: 45, 36, 33 and 21, respectively, for the same four contrasts. In TOLENTINO-2022, no miRNA reached significance (padj < 0.10) in any of the three contrasts under the mixed statistical design we used.

### 3.6 Predicted target landscape

The current release contains 1,773 predicted miRNA-target interactions generated with psRNATarget, spanning 764 distinct target loci; all 99 catalog entries (100%) are linked to at least one predicted target. The protein-protein interaction (PPI) network retrieved for these target loci, together with its associated functional enrichment, is shown in Figure 5.

**Figure 5.**
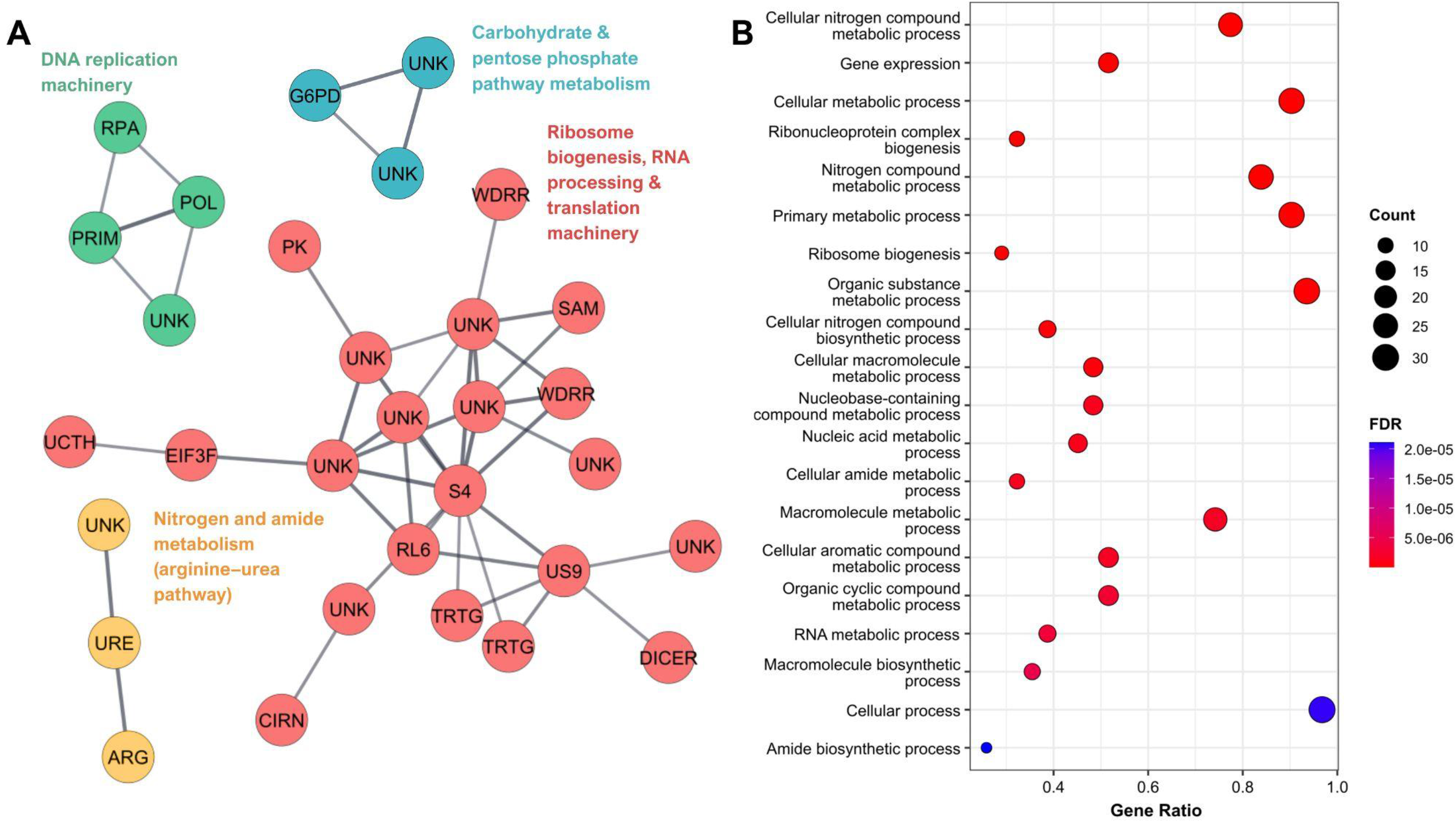
Functional characterization of *E. grandis* miRNA target loci identified in EMA. **(A)** Protein-protein interaction network retrieved via the STRING v12.5 database at a high-confidence threshold (≥ 0.700), with no additional interactors and disconnected nodes removed. (B) Dot plot of the 20 most significantly enriched GO Biological Process terms for the same protein set, ranked by FDR; dot size denotes the number of proteins annotated to each term, and color denotes FDR.

At this confidence threshold, the network (Figure 5A) comprises 31 connected target proteins and 44 edges, organized into 4 self-contained clusters. The ribosome biogenesis, RNA processing and translation machinery cluster is the largest, comprising a Dicer-like RNase III protein, an S4 RNA-binding protein, ribosomal proteins L6 and uS9, a SAM-dependent rRNA methyltransferase of the RsmB/NOP family, two Tr-type G domain translation elongation factors, eukaryotic translation initiation factor 3 subunit F (eIF3F), two WD-repeat proteins, a Cir_N domain protein, a ubiquitin C-terminal hydrolase, a protein kinase, and several uncharacterized proteins. The DNA replication machinery cluster comprises a DNA polymerase, a DNA primase large subunit, and a Replication Protein A subunit, together with an additional uncharacterized protein. The nitrogen and amide metabolism (arginine-urea pathway) cluster comprises arginase, urease, and an uncharacterized protein. The carbohydrate and pentose phosphate pathway metabolism cluster comprises glucose-6-phosphate dehydrogenase and two additional uncharacterized proteins.

Functional enrichment of this protein set (Figure 5B) is dominated, in both significance and gene count, by a cascade of increasingly broad, hierarchically nested GO Biological Process terms centered on nitrogen-compound and primary metabolism (GO:0034641, Cellular nitrogen compound metabolic process, 24 of 31 proteins, FDR = 3.15e-14; GO:0044237, Cellular metabolic process, 28 proteins; GO:0044238, Primary metabolic process, 28 proteins; GO:0071704, Organic substance metabolic process, 29 proteins).

### 3.7 Functional sections of the EMA web interface

The EMA web interface is organized into successive levels of information, from global database summaries to individual miRNA evidence records, as illustrated in Figure 6.

**Figure 6.**
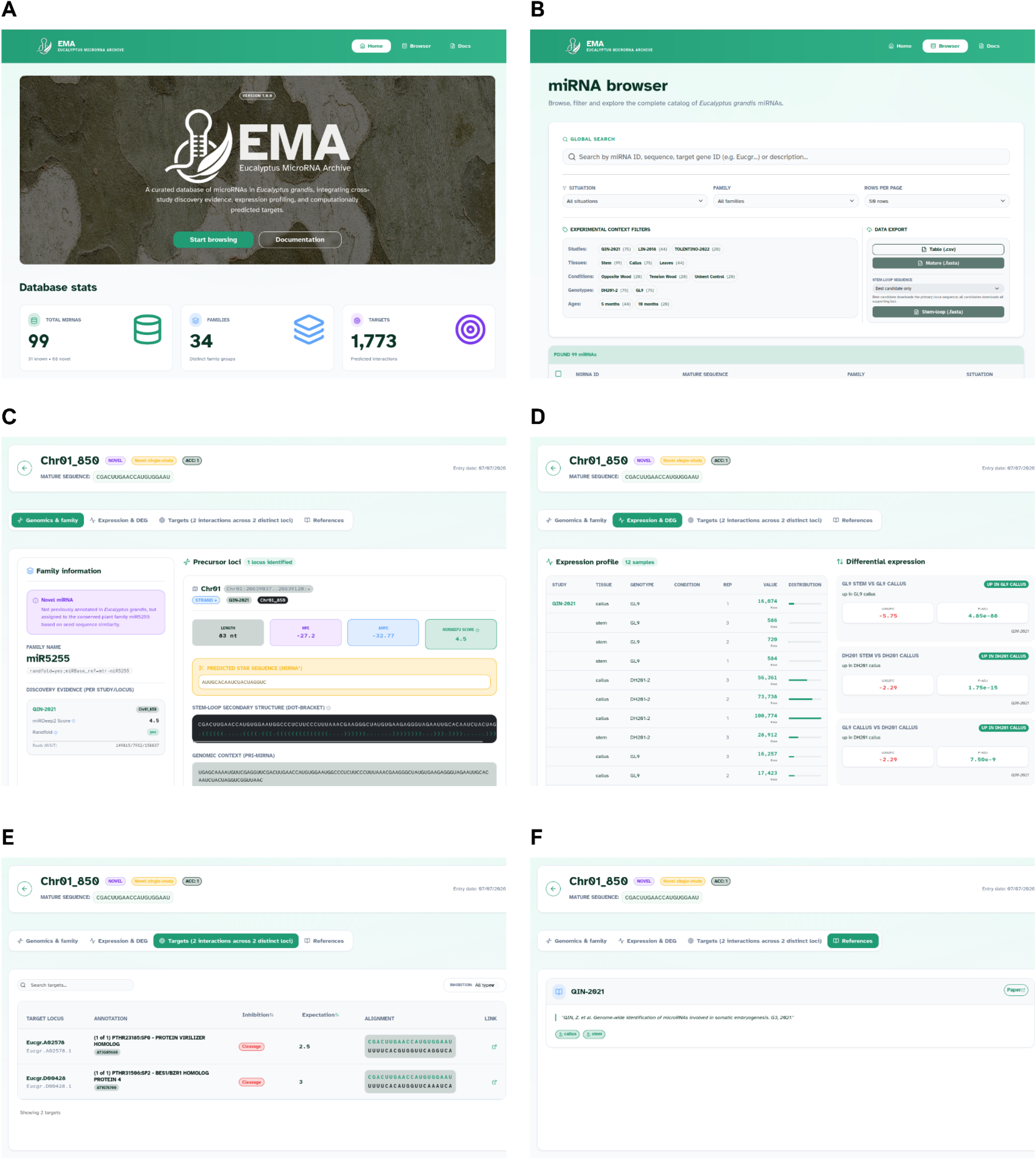
Main sections of the EMA web interface: (A) home page; (B) data browser; (C) miRNA entry overview; (D) miRNA quantification data; (E) miRNA targets; and (F) entry references and sample information.

The landing page (Figure 6A) provides an overview of the current release. It also provides direct access to the data browser (Figure 6B), which enables users to search the catalog by miRNA identifier, mature sequence, target gene identifier, or target description, and to filter entries by biological context and experimental metadata, including study, tissue, condition, genotype, and plant age. The miRNA detail page (Figure 6C) links each canonical EMA entry to its supporting evidence, structural features, and confidence tier (known/reference-supported, novel/multi-study replicated, or novel/single-study), while displaying the accession, mature sequence, family assignment, study-specific discovery evidence, and precursor coordinates and secondary structure. The quantification section (Figure 6D) provides expression profiles across sampled conditions and differential expression results for statistically significant comparisons. The targets section (Figure 6E) lists predicted miRNA targets together with their functional annotations, enabling users to explore the potential biological roles associated with each miRNA. Finally, the references and samples section (Figure 6F) links each entry to the studies supporting its annotation and provides detailed information on the experimental samples, including the biological and experimental contexts in which the miRNA was identified or quantified.

## 4 Discussion

### 4.1 EMA addresses a fragmented annotation landscape

*E. grandis* miRNAs have been reported several times over the past decade, with substantial variation in reported numbers. The original genome annotation described 206 loci in 36 families [5]. [6] reported 95 conserved and 193 novel miRNAs in a degradome-validated survey [6]. [4] reported 40 known and 8 novel miRNAs in a developmental study, though library composition was dominated by t/rRNA contamination and discovery relied on a single non-replicated library, a profile common to several pre-2018 efforts and one that the annotation criteria adopted here are specifically designed to screen out [4]. [7], one of EMA’s three source datasets, reported 179 novel miRNAs in 41 families in leaf tissue and 257 novel miRNAs in 61 families in stem tissue [7]. The remaining two source datasets integrated here, [8] and [9], reported their own respective miRNA sets for somatic embryogenesis and tension wood [8,9]. These original studies also differ markedly in scope and method, reported family counts range from 36 to 61, filtering criteria, expression data, and genomic coordinate conventions are not directly comparable across studies, and in some cases the underlying raw sequencing data are no longer accessible. EMA addresses this fragmentation directly, that is, by requiring public raw-data availability as an inclusion criterion and applying a single, uniform annotation and confidence-tiering framework across all integrated studies, replacing study-specific, non-comparable criteria with one reproducible standard applied consistently to every entry in the catalog.

PmiREN [13], a multi-species resource spanning approximately 38,000 miRNA loci across 179 plant species, currently lists 82 miRNA loci in 58 families for *E. grandis*, distributed across 6 clusters, with no sRNA-seq or PARE-seq datasets and no syntenic blocks recorded for this species in its metadata. This entry is smaller in absolute count than the 99-entry EMA catalog and, as a species-level snapshot, does not preserve relevant metadata such as study of origin, sample-level read support, or condition-specific differential expression. EMA’s contribution is a locus- and evidence-level reconciliation across three biologically defined contexts (vegetative tissue, somatic embryogenesis, tension wood) that keeps every entry traceable to its supporting dataset, closing a gap that neither individual primary studies nor pan-species databases currently fill for this species.

### 4.2 Cross-study reproducibility justifies an evidence-based catalog

The dominant UpSet intersection (Figure 2A) reflects reproducibility within a single deeply replicated study (QIN-2021’s four sample subgroups) rather than agreement between studies: only 9 of 99 miRNAs (9.1%) are supported by all three studies, and 68 (68.7%) rest on single-study evidence. The discovery-score comparison (Figure 2C) adds a related observation. TOLENTINO-2022’s high mean score is driven by a small evidence set (n = 27) and a few extreme values, while its median is the lowest of the three studies, whereas QIN-2021 combines the highest median with the broadest spread, consistent with deeper discovery power rather than outlier inflation. Structural evidence follows the same pattern: single-study novel candidates show greater heterogeneity in precursor length and MFE (Figure 3B) than reference-supported or multi-study replicated entries. These patterns justify EMA’s three-tier confidence system (known-reference-supported, novel multi-study replicated, novel single-study) instead of a binary known/novel split, and indicate where future validation effort should be directed.

### 4.3 Limitations and validation priorities

The curation applied throughout this work concerns miRNA locus annotation: the discovery, structural filtering, and cross-study reconciliation processes described in section 2. It does not extend to the predicted target interactions reported, which remain computational outputs, not experimentally validated relationships. Cross-referencing EMA’s target predictions, particularly for Egr-miR482b, against the degradome- and PARE-supported target set independently reported for *E. grandis* by Pappas [6] would constitute a direct, low-cost step toward experimentally supporting a subset of these predictions in a future release. Multi-species resources such as PmiREN incorporate PARE/degradome-based target validation where such data exists for a given species [12]; the absence of this layer for *E. grandis*, in both PmiREN’s own entry and in the present release, is a shared gap that the [6] dataset is well positioned to help close.

## 5 Conclusion

EMA consolidates three independently generated small RNA sequencing datasets into a single, locus-resolved catalog of 99 curated miRNAs for *E. grandis*, spanning vegetative development, somatic embryogenesis, and mechanically induced wood formation. Unlike prior single-study annotations or species-wide snapshots such as PmiREN, EMA preserves study-of-origin, sample-level evidence, and condition-specific expression for every entry, organized under an explicit three-tier confidence system that distinguishes reference-supported, multi-study replicated, and single-study candidates. This structure gives researchers a transparent basis for prioritizing candidates for further study, rather than treating the catalog as a uniform list of equally supported loci. The underlying curation pipeline, confidence-tiering framework, and relational schema were built independently of any species-specific assumption, and the same architecture can directly accommodate future *E. grandis* datasets, as well as sequencing data from related *Eucalyptus* and Myrtaceae species, without structural modification. Together with its public web interface and open codebase, EMA therefore serves not only as a resource for *E. grandis,* but as a reusable, extensible framework positioned to scale with the growing volume of small RNA sequencing data across woody, non-model plant species.

## Acknowledgements

The authors thank the Venancio Group, Laboratory of Chemistry and Function of Peptides and Proteins (LQPFPP), Universidade Estadual do Norte Fluminense Darcy Ribeiro (UENF), for hosting the EMA web dashboard.

## Data availability statement

The curated SQLite database, backend schema, exported catalog and evidence tables, network construction and functional annotation scripts, and pipeline execution scripts and logs are available at the following GitHub repository: github.com/jvtarss/ema-2026.

## REFERENCES

[1] M.J. Axtell, B.C. Meyers, Revisiting Criteria for Plant MicroRNA Annotation in the Era of Big Data, Plant Cell 30 (2018) 272–284. 10.1105/tpc.17.00851.

[2] L. Fang, Y. Wang, MicroRNAs in Woody Plants, Front. Plant Sci. 12 (2021) 686831. 10.3389/fpls.2021.686831.

[3] A.M. Belkozhayev, A. Abaildayev, B.D. Kossalbayev, A. Kerimkulova, D.K. Kadirshe, G. Toleutay, The Role of MicroRNA-Based Strategies in Optimizing Plant Biomass Composition for Bio-Based Packaging Materials, Plants 14 (2025) 2905. 10.3390/plants14182905.

[4] A. Levy, D. Szwerdszarf, M. Abu-Abied, I. Mordehaev, Y. Yaniv, J. Riov, T. Arazi, E. Sadot, Profiling microRNAs in Eucalyptus grandis reveals no mutual relationship between alterations in miR156 and miR172 expression and adventitious root induction during development, BMC Genomics 15 (2014) 524. 10.1186/1471-2164-15-524.

[5] A.A. Myburg, D. Grattapaglia, G.A. Tuskan, U. Hellsten, R.D. Hayes, J. Grimwood, J. Jenkins, E. Lindquist, H. Tice, D. Bauer, D.M. Goodstein, I. Dubchak, A. Poliakov, E. Mizrachi, A.R.K. Kullan, S.G. Hussey, D. Pinard, K. Van Der Merwe, P. Singh, I. Van Jaarsveld, O.B. Silva-Junior, R.C. Togawa, M.R. Pappas, D.A. Faria, C.P. Sansaloni, C.D. Petroli, X. Yang, P. Ranjan, T.J. Tschaplinski, C.-Y. Ye, T. Li, L. Sterck, K. Vanneste, F. Murat, M. Soler, H.S. Clemente, N. Saidi, H. Cassan-Wang, C. Dunand, C.A. Hefer, E. Bornberg-Bauer, A.R. Kersting, K. Vining, V. Amarasinghe, M. Ranik, S. Naithani, J. Elser, A.E. Boyd, A. Liston, J.W. Spatafora, P. Dharmwardhana, R. Raja, C. Sullivan, E. Romanel, M. Alves-Ferreira, C. Külheim, W. Foley, V. Carocha, J. Paiva, D. Kudrna, S.H. Brommonschenkel, G. Pasquali, M. Byrne, P. Rigault, J. Tibbits, A. Spokevicius, R.C. Jones, D.A. Steane, R.E. Vaillancourt, B.M. Potts, F. Joubert, K. Barry, G.J. Pappas, S.H. Strauss, P. Jaiswal, J. Grima-Pettenati, J. Salse, Y. Van De Peer, D.S. Rokhsar, J. Schmutz, The genome of Eucalyptus grandis, Nature 510 (2014) 356–362. 10.1038/nature13308.

[6] M.D.C.R. Pappas, G.J. Pappas, D. Grattapaglia, Genome-wide discovery and validation of Eucalyptus small RNAs reveals variable patterns of conservation and diversity across species of Myrtaceae, BMC Genomics 16 (2015) 1113. 10.1186/s12864-015-2322-6.

[7] Z. Lin, Q. Li, Q. Yin, J. Wang, B. Zhang, S. Gan, A.-M. Wu, Identification of novel miRNAs and their target genes in Eucalyptus grandis, Tree Genet. Genomes 14 (2018) 60. 10.1007/s11295-018-1273-x.

[8] Z. Qin, J. Li, Y. Zhang, Y. Xiao, X. Zhang, L. Zhong, H. Liu, B. Chen, Genome-wide identification of microRNAs involved in the somatic embryogenesis of Eucalyptus, G3 GenesGenomesGenetics 11 (2021) jkab070. 10.1093/g3journal/jkab070.

[9] F.T. Tolentino, A.A. Vasconcelos, U.R. Souza, G.A.G. Pereira, M.F. Carazolle, P. Mazzafera, Identification of microRNAs and their expression profiles on tension and opposite wood of Eucalyptus, Theor. Exp. Plant Physiol. 34 (2022) 485–500. 10.1007/s40626-022-00259-9.

[10] B. Alptekin, B.A. Akpinar, H. Budak, A Comprehensive Prescription for Plant miRNA Identification, Front. Plant Sci. 7 (2017). 10.3389/fpls.2016.02058.

[11] R. Yang, H. Zhang, Application of Artificial Intelligence Technology in Plant MicroRNA Research: Progress, Challenges, and Prospects, Int. J. Mol. Sci. 26 (2025) 11854. 10.3390/ijms262411854.

[12] Z. Guo, Z. Kuang, Y. Wang, Y. Zhao, Y. Tao, C. Cheng, J. Yang, X. Lu, C. Hao, T. Wang, X. Cao, J. Wei, L. Li, X. Yang, PmiREN: a comprehensive encyclopedia of plant miRNAs, Nucleic Acids Res. 48 (2020) D1114–D1121. 10.1093/nar/gkz894.

[13] Z. Guo, Z. Kuang, Y. Zhao, Y. Deng, H. He, M. Wan, Y. Tao, D. Wang, J. Wei, L. Li, X. Yang, PmiREN2.0: from data annotation to functional exploration of plant microRNAs, Nucleic Acids Res. 50 (2022) D1475–D1482. 10.1093/nar/gkab811.

[14] B.C. Meyers, M.J. Axtell, B. Bartel, D.P. Bartel, D. Baulcombe, J.L. Bowman, X. Cao, J.C. Carrington, X. Chen, P.J. Green, S. Griffiths-Jones, S.E. Jacobsen, A.C. Mallory, R.A. Martienssen, R.S. Poethig, Y. Qi, H. Vaucheret, O. Voinnet, Y. Watanabe, D. Weigel, J.-K. Zhu, Criteria for Annotation of Plant MicroRNAs, Plant Cell 20 (2008) 3186–3190. 10.1105/tpc.108.064311.

[15] A.A. Myburg, D. Grattapaglia, G.A. Tuskan, U. Hellsten, R.D. Hayes, J. Grimwood, J. Jenkins, E. Lindquist, H. Tice, D. Bauer, D.M. Goodstein, I. Dubchak, A. Poliakov, E. Mizrachi, A.R.K. Kullan, S.G. Hussey, D. Pinard, K. Van Der Merwe, P. Singh, I. Van Jaarsveld, O.B. Silva-Junior, R.C. Togawa, M.R. Pappas, D.A. Faria, C.P. Sansaloni, C.D. Petroli, X. Yang, P. Ranjan, T.J. Tschaplinski, C.-Y. Ye, T. Li, L. Sterck, K. Vanneste, F. Murat, M. Soler, H.S. Clemente, N. Saidi, H. Cassan-Wang, C. Dunand, C.A. Hefer, E. Bornberg-Bauer, A.R. Kersting, K. Vining, V. Amarasinghe, M. Ranik, S. Naithani, J. Elser, A.E. Boyd, A. Liston, J.W. Spatafora, P. Dharmwardhana, R. Raja, C. Sullivan, E. Romanel, M. Alves-Ferreira, C. Külheim, W. Foley, V. Carocha, J. Paiva, D. Kudrna, S.H. Brommonschenkel, G. Pasquali, M. Byrne, P. Rigault, J. Tibbits, A. Spokevicius, R.C. Jones, D.A. Steane, R.E. Vaillancourt, B.M. Potts, F. Joubert, K. Barry, G.J. Pappas, S.H. Strauss, P. Jaiswal, J. Grima-Pettenati, J. Salse, Y. Van De Peer, D.S. Rokhsar, J. Schmutz, The genome of Eucalyptus grandis, Nature 510 (2014) 356–362. 10.1038/nature13308.

[16] S. Chen, Y. Zhou, Y. Chen, J. Gu, *fastp* : an ultra-fast all-in-one FASTQ preprocessor, (2018). 10.1101/274100.

[17] Y. Fu, P.-H. Wu, T. Beane, P.D. Zamore, Z. Weng, Elimination of PCR duplicates in RNA-seq and small RNA-seq using unique molecular identifiers, BMC Genomics 19 (2018) 531. 10.1186/s12864-018-4933-1.

[18] M.R. Friedländer, S.D. Mackowiak, N. Li, W. Chen, N. Rajewsky, miRDeep2 accurately identifies known and hundreds of novel microRNA genes in seven animal clades, Nucleic Acids Res. 40 (2012) 37–52. 10.1093/nar/gkr688.

[19] K. Pawlina-Tyszko, T. Szmatoła, Benchmarking of bioinformatics tools for NGS-based microRNA profiling with RT-qPCR method, Funct. Integr. Genomics 23 (2023) 347. 10.1007/s10142-023-01276-w.

[20] A. Kozomara, M. Birgaoanu, S. Griffiths-Jones, miRBase: from microRNA sequences to function, Nucleic Acids Res. 47 (2019) D155–D162. 10.1093/nar/gky1141.

[21] E. Bonnet, J. Wuyts, P. Rouzé, Y. Van De Peer, Evidence that microRNA precursors, unlike other non-coding RNAs, have lower folding free energies than random sequences, Bioinformatics 20 (2004) 2911–2917. 10.1093/bioinformatics/bth374.

[22] G. Deschamps-Francoeur, J. Simoneau, M.S. Scott, Handling multi-mapped reads in RNA-seq, Comput. Struct. Biotechnol. J. 18 (2020) 1569–1576. 10.1016/j.csbj.2020.06.014.

[23] M. Zytnicki, C. Gaspin, srnaMapper: an optimal mapping tool for sRNA-Seq reads, BMC Bioinformatics 23 (2022) 495. 10.1186/s12859-022-05048-4.

[24] K. Ciechanowska, M. Pokornowska, A. Kurzyńska-Kokorniak, Genetic Insight into the Domain Structure and Functions of Dicer-Type Ribonucleases, Int. J. Mol. Sci. 22 (2021) 616. 10.3390/ijms22020616.

[25] P. Li, Y. Piao, H.S. Shon, K.H. Ryu, Comparing the normalization methods for the differential analysis of Illumina high-throughput RNA-Seq data, BMC Bioinformatics 16 (2015) 347. 10.1186/s12859-015-0778-7.

[26] M.I. Love, W. Huber, S. Anders, Moderated estimation of fold change and dispersion for RNA-seq data with DESeq2, Genome Biol. 15 (2014) 550. 10.1186/s13059-014-0550-8.

[27] Z. Zhang, D. Tian, J. Wang, P. He, X. Yang, L. Jiang, Impacts of tree age and external morphological traits on longitudinal variation of stem wood density for mixed forests, For. Ecol. Manag. 598 (2025) 123255. 10.1016/j.foreco.2025.123255.

[28] X. Dai, Z. Zhuang, P.X. Zhao, psRNATarget: a plant small RNA target analysis server (2017 release), Nucleic Acids Res. 46 (2018) W49–W54. 10.1093/nar/gky316.

[29] D. Szklarczyk, K. Nastou, M. Koutrouli, R. Kirsch, F. Mehryary, R. Hachilif, D. Hu, M.E. Peluso, Q. Huang, T. Fang, N.T. Doncheva, S. Pyysalo, P. Bork, L.J. Jensen, C. von Mering, The STRING database in 2025: protein networks with directionality of regulation, Nucleic Acids Res. 53 (2025) D730–D737. 10.1093/nar/gkae1113.

[30] P. Shannon, A. Markiel, O. Ozier, N.S. Baliga, J.T. Wang, D. Ramage, N. Amin, B. Schwikowski, T. Ideker, Cytoscape: A Software Environment for Integrated Models of Biomolecular Interaction Networks, Genome Res. 13 (2003) 2498–2504. 10.1101/gr.1239303.

[31] H. Wickham, ggplot2, Springer International Publishing, Cham, 2016. 10.1007/978-3-319-24277-4.

[32] M. Bayer, SQLAlchemy, in: A. Brown, G. Wilson (Eds.), Archit. Open Source Appl. Vol. II Struct. Scale Few More Fearless Hacks, aosabook.org, Mountain View, 2012. http://aosabook.org/en/sqlalchemy.html.

[33] S. Ramírez, FastAPI, (2018). https://github.com/fastapi/fastapi.

[34] S. Colvin, E. Jolibois, H. Ramezani, A. Garcia Badaracco, T. Dorsey, D. Montague, S. Matveenko, M. Trylesinski, S. Runkle, D. Hewitt, A. Hall, V. Plot, Pydantic validation, (2026). https://github.com/pydantic/pydantic.

